# Juvenile agile frogs spatially avoid ranavirus-infected conspecifics, but do not show generalized social distancing or self-isolation

**DOI:** 10.1101/2024.04.10.588850

**Authors:** Dávid Herczeg, Gergely Horváth, Veronika Bókony, Gábor Herczeg, Andrea Kásler, Dóra Holly, Zsanett Mikó, Nikolett Ujhegyi, János Ujszegi, Tibor Papp, Attila Hettyey

## Abstract

Exposure to contagious pathogens can result in behavioural changes, which can alter the spread of infectious diseases. Healthy individuals can express generalized social distancing or actively avoid the sources of infection, while infected individuals can show passive or active self-isolation. Amphibians are globally threatened by serious contagious diseases, yet their behavioural responses to infections are very scarcely known. We studied behavioural changes in agile frog (*Rana dalmatina*) juveniles upon exposure to a *Ranavirus* (*Rv*). We performed classic choice tests in chambers containing a conspecific infected with *Rv* in one end compartment and a non-infected conspecific in the other end, with an *Rv*-infected or non-infected focal individual in the central compartment. We found that both non-infected and *Rv*- infected focal individuals spatially avoided infected conspecifics, while there were no signs of generalized social distancing, nor self-isolation. Spatial avoidance of infected conspecifics may effectively hinder disease transmission. On the other hand, the absence of self-isolation by infected individuals may facilitate it. Our finding that infected individuals spent more time near the non-infected than infected conspecifics suggests that the strong behavioural drive to avoid infected conspecifics may not be silenced by infection, possibly to prevent secondary infections. The observation that infected focal individuals did not spend more time near conspecifics than non-infected focals renders it unlikely that the pathogen manipulated host behaviour to aid disease spread. More research is urgently needed to understand under what circumstances behavioural responses can help amphibians cope with infections, and how that affects disease dynamics in natural populations.

**GRAPHICAL ABSTRACT:** 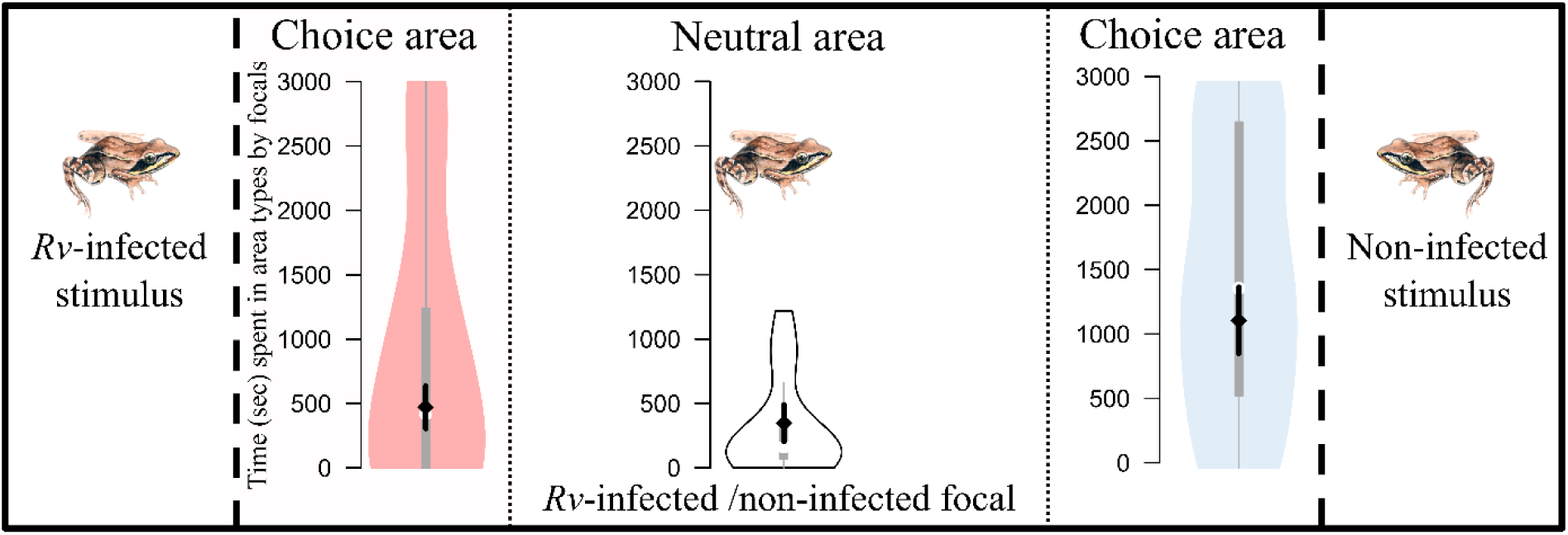

## INTRODUCTION

Detailed knowledge regarding pathogen-mediated behavioural changes in hosts is important for understanding the spread of infectious diseases (Stockmaier et al. 2021) and for the design and implementation of informed and effective mitigation interventions. This is because not only an effective immune system, but also behavioural mechanisms can contribute to the anti-pathogen defence of animals, and influence disease dynamics in natural populations. Healthy individuals can decrease the chances of becoming infected *via* generalized social distancing, i.e. by reducing the frequency or duration of social interactions non-selectively with all social partners (Stroeymeyt et al. 2014). In addition, if individuals are capable of recognizing signs of infection, or more generally, contamination, they can reduce transmission risk by displaying avoidance behaviours, where they spatially avoid individuals carrying the pathogen (Behringer et al. 2006; Paciência et al. 2019), locations of elevated risk of infection (Kiesecker & Skelly 2000; Sarabian et al. 2018; Weinstein et al. 2018), food sources that are contaminated (Moleón et al. 2017; Amoroso et al. 2019) or encounters with the parasites or their vectors (Behringer et al. 2018). Furthermore, infected individuals are often characterized by sickness behaviours, where they become apathetic, inactive, and socially disinterested due to tissue damage, the side effects of mounting an immune response, and elevated energy needs of fighting the infection, which in turn lower the chance of passing on the disease. This leads to passive self-isolation. Passive self-isolation occurs when an infected individual reduces contact with conspecifics directly *via* social disinterest (in social animals) or indirectly *via* decreased movement activity, i.e. lethargy (Kazlauskas et al. 2016; Geffre et al. 2020; Stockmaier et al. 2020). On the other hand, self-isolation can also happen actively, whereby infected individuals maintain their activity but direct it away from social partners (Heinze & Walter 2010; Bos et al. 2012; Stroeymeyt et al. 2018). All types of the above behavioural changes have been documented to occur in humans and eusocial insects. However, only avoidance behaviours and passive self-isolation appear to be widespread throughout the animal kingdom, whereas reports of generalized social distancing and active self-isolation are extremely rarely documented (Heinze & Walter 2010; Bos et al. 2012; Stroeymeyt et al. 2018). Finally, not all behavioural responses are adaptive for hosts, as some pathogens can manipulate host behaviour to increase the chances of transmission (Hafer 2016).

Amphibians are the most threatened class of vertebrates (Luedtke et al. 2023). During recent decades, their populations suffered sharp declines (Wake & Vredenburg 2008) resulting in 40.7 % of species being globally threatened (Luedtke et al. 2023). The mechanisms behind these declines are complex (Collins 2010), but primarily driven by climate change, habitat loss and infectious diseases (Scheele et al. 2019; Fisher & Garner 2020; Luedtke et al. 2023). Nonetheless, behavioural changes following exposure to a pathogen were seldom studied in amphibians. The handful of studies that did so (Kiesecker et al. 1999; McMahon et al. 2014; McMahon et al. 2021; Le Sage et al. 2022) reported that healthy individuals avoided infected conspecifics or contaminated surfaces and did not show generalized social distancing. Infected amphibians can show anorexia and lethargy (Gray et al. 2009; Landsberg et al. 2013), but whether these result in passive self-isolation and whether amphibians display active self-isolation is not yet known.

Ranaviruses (family *Iridoviridae*; hereafter *Rv*) are a group of aquatic pathogens with a wide host range among ectothermic vertebrates, including amphibians (Duffus et al. 2015). Epidemics caused by *Rv* have led to mass mortality events across Europe, Asia and the Americas (Gray et al. 2009), resulting in local extinctions of amphibians (Kik et al. 2011; Price et al. 2014). Ranavirus*-*associated mortality can reach 100 % after metamorphosis in some amphibian species (Schock et al. 2008; Earl et al. 2016), but interspecific variation in susceptibility is high (Gray et al. 2009). Transmission mainly occurs by direct contact between individuals, but can also happen *via* ingestion of infected tissue or exposure to water or a wet substrate containing virions (Brunner et al. 2015). Field data and experimental infection studies on amphibians described a behavioural change following *Rv*-infection with individuals displaying lethargy as a sign of passive self-isolation (Gray et al. 2009). The only study testing for avoidance behaviour in metamorphosed amphibians concerning *Rv* found that healthy wood frogs (*Rana sylvatica*) kept greater distances from *Rv*-infected than from healthy conspecifics, which could be interpreted as avoidance behaviour (Le Sage et al. 2022). Furthermore, the observed distance kept to infected individuals was positively correlated with infection intensity (Le Sage et al. 2022). However, the animals tested in a given trial were entered together into an open arena, where they could move around without restriction. Consequently, their behaviour could not be considered independent from each other and, partly as a consequence, the different elements of behavioural changes (i.e., generalized social distancing, avoidance behaviour and passive/active self-isolation) could not be separated unambiguously.

In the present paper, we examined how the behaviour of juvenile agile frogs *Rana dalmatina* changes when facing *Rv* (Herczeg et al. 2023). To this end, we ran classic choice tests, where the focal and the stimulus individuals were spatially separated from each other by transparent and perforated walls. This setup allowed for the exchange of visual and chemical information and allowed the focal individual to adjust its behaviour to its internal state and its environment while not under threat of being approached by potentially infectious conspecifics. This allowed us to discern with unprecedented detail the various forms of behavioural changes in amphibians following exposure to a pathogen. We hypothesized that the behaviour of agile frogs depended on their infection status and on that of their conspecifics in their immediate environment. We predicted that if healthy froglets were able to sense the presence of the pathogen, they would either avoid all conspecifics (generalized social distancing) or spatially avoid *Rv*-infected conspecifics (avoidance behaviour), while infected individuals would show decreased movement activity (passive self-isolation *via* lethargy) or general conspecific avoidance (active self-isolation). In case *Rv* can manipulate hosts to enhance its chances of transmission, we expected to observe infected focal individuals spend more time near conspecifics than non-infected focals. Furthermore, if hosts can discern between infected and non-infected conspecifics and *Rv* evolved highly fine-tuned manipulation, we may see infected focals spending more time close to non-infected conspecifics, than non-infected focals. Finally, we expected the magnitude of behavioural changes to be positively correlated to *Rv* infection intensity.

## MATERIALS AND METHODS

### Collection and rearing of animals

In April 2022, we collected 200 eggs from each of eight freshly laid agile frog egg clutches (hereafter sibgroups) from a temporary pond (Ilona-tó, 47.71326 N, 19.04050 E), located in the hilly woodland of the Pilis-Visegrádi Mountains in Hungary. We transported eggs to the Júliannamajor Experimental Station of the Plant Protection Institute, HUN-REN Centre for Agricultural Research in Budapest, Hungary. For a schematic timeline of the experiment see Fig. 1A. Briefly, until development stage 25 (Gosner 1960) we kept each sibgroup separately in a plastic container (24 × 16 × 13 cm) filled with 1 L reconstituted soft water (RSW) (USEPA 2002) in the laboratory at a constant temperature of 19 °C and a 12:12 h light:dark cycle. Meanwhile, we set up 16 outdoor mesocosms (80 × 55 × 36 cm) by filling them with 130 L of aged tap water and adding 40 g dried beech *Fagus sylvatica* leaves to provide nutrients and shelter to tadpoles. To enhance algal growth and start up a self-sustaining ecosystem, we inoculated mesocosms with 1 L of pond water containing bacterio-, phyto-, and zooplankton. When tadpoles reached development stage 25, we haphazardly selected two groups of 50 healthy-looking tadpoles from each sibgroup and placed them into the mesocosms. When the first individual reached development stage 42 (emergence of forelimbs, i.e., the start of metamorphosis), we removed leaves from all mesocosms and monitored them daily for individuals reaching development stage 42. If we found one, we placed it into a semi-transparent, 45 L plastic box (56 × 39 × 28 cm; i.e., one box for each mesocosm) covered with a perforated lid and placed into a shady outdoor area. Plastic boxes contained 0.5 L of aged tap water and were slightly tilted to provide both water cover as well as a dry area. After development stage 46 (i.e., the end of metamorphosis), we re-located juveniles to the laboratory, where we kept them individually in 1.5 L plastic boxes covered with a perforated lid and containing moistened paper towels as substrate and a piece of egg carton to provide shelter. Twice a week, we sprinkled boxes with RSW to maintain humidity and fed metamorphs *ad libitum* with small crickets (*Acheta domesticus*, instar stage 1–2) until the start of the assays.

**Figure 1:**
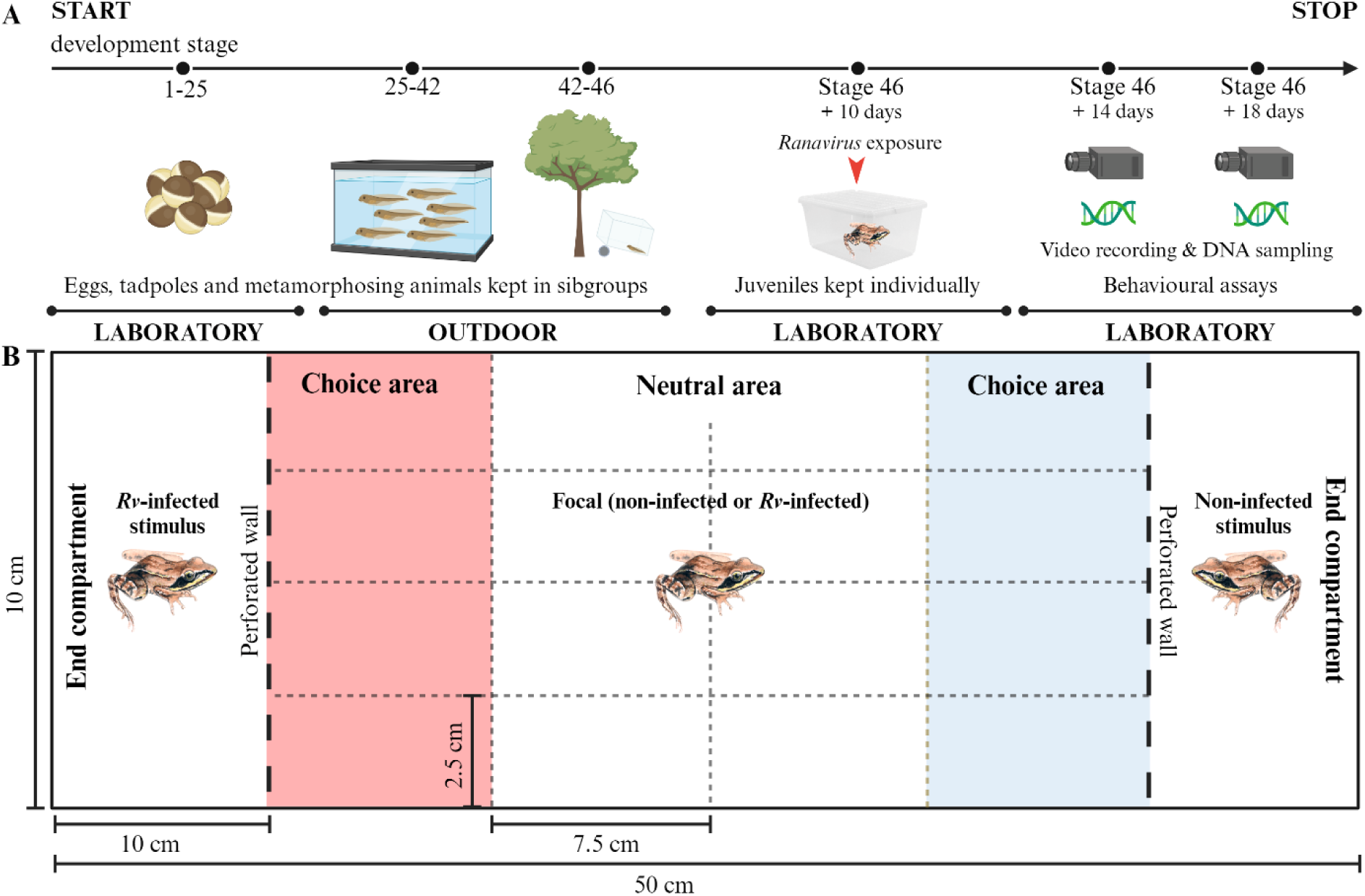
A schematic timeline of the experiment (A), the time shown as the development stage according to (Gosner 1960), and the compartments of the choice chambers as viewed from above (B).

Ten days after completion of metamorphosis we exposed half of the agile frogs (sibgroups represented equally and animals assigned to treatments randomly) to *Rv* (Frog Virus 3; ATCC No. VR-567; Fig. 1A). For the maintenance of *Rv* cultures see more details in Herczeg et al. (2023). We placed each froglet in a 6 cm diameter Petri dish filled with 8 ml RSW and containing 5×10^5^ plaque forming unit (pfu) × ml^-1^ of the virus. In the case of controls, we added the same quantity of sham extract (only the medium nutrient medium with 2 % fetal bovine serum without the virus) to the Petri dishes. Five hours later, we placed the juveniles back into their original boxes.

### Behavioural assays

We used 16 rectangular choice chambers (Fig. 1B) with end compartments on both ends. The side walls of the choice chambers were non-transparent. The end compartments were separated from the central compartment with perforated and transparent Plexiglas walls (3 mm in thickness, with nine 2.5 mm wide slits running from the bottom to close to the top), allowing for the exchange of both visual and chemical cues, but preventing physical contact and pathogen transmission between neighbouring compartments. We did not provide any objects as shelters to avoid interference with visual and chemical cues. We covered the bottom of all three compartments of the choice chambers with wet paper towels to avoid the desiccation of froglets during the trials and placed a sliding Plexiglas lid on top to prevent juveniles from escaping. After each trial, we discarded the paper towels, disinfected the choice chambers by spraying with 70 % ethanol, wiped chambers to remove excess alcohol and allowed at least 1 hour for complete drying. Finally, we lined compartments with new paper towels and wet them with RSW.

We performed behavioural trials 14 or 18 days after metamorphosis (i.e., 4 or 8 days after exposure to *Rv*) using randomly created triads of individuals, where each individual was used only once. We performed trials at two-time points because we had no previous information about the temporal dimension of disease progression in *Rv*-infected agile frog juveniles. We started trials by manually placing focal individuals into the middle of the central compartments. First, we let focal individuals acclimatize and explore the arena in the initial part for 30 minutes (also to assess the baseline behaviour of focals). Subsequently, we placed one infected and one non-infected stimulus animal into the end compartments, repositioned the focal individuals back to the centre of the central compartment and ran trials for another 60 minutes (to assess social and sickness behaviour in focals). We recorded the movement of animals using HD video cameras (Canon Legria Hf R66) attached to tripods and positioned above the choice chambers. The 16 choice chambers were randomly divided into two groups, one for assessing the behaviour of healthy focal individuals and the other one for assessing that of infected focals. We also randomised for each chamber which end compartment will house healthy *vs.* infected stimulus individuals. This way, we reached maximum spatial randomisation, while minimizing the possibility of cross-contamination and avoiding hampering the effects of cues potentially remaining in the end compartments despite cleaning. At the end of trials, we measured body mass to the nearest 0.01 g using a laboratory scale (Ohaus PA114), euthanized individuals with the ‘cooling then freezing’ method (Shine et al. 2015) and conserved them in 3 ml of 96 % ethanol until further molecular analysis.

### Ranavirus detection with qPCR

To assess *Rv* infection intensity, we homogenised liver tissues with a disposable pellet mixer (VWR, catalogue no. 47747-370) and extracted the *Rv* DNA with Wizard Genomic DNA Purification Kit (Promega, Madison, Wisconsin, USA) according to the manufacturer’s protocol. We stored extracted DNA at -20 °C until further analyses. We estimated *Rv* infection intensity by targeting the major capsid protein (MCP) gene of the viral genome following standard amplification methodologies (Stilwell et al. 2018). We ran samples in duplicate, and when the result was equivocal, we repeated reactions in duplicate. If we obtained an equivocal result again, we considered the sample positive (Stilwell et al. 2018). Reactions were run on a BioRad CFX96 Touch Real-Time PCR System. For further details see Herczeg et al. (2023).

### Video analysis

During the trials, the movement of focal individuals was coded as a state event (i.e., a behaviour with duration) and analysed by a single observer with the BORIS event logging software (Friard & Gamba 2016). We trimmed the beginning and the end of each recording (5 min) to exclude any possible disturbance caused by the experimenters exiting and entering the experimental room. This resulted in the analysis of 20 minutes of baseline behaviour and 50 minutes of observing social and sickness behaviours following the addition of stimulus individuals. We divided the central compartment of the choice chambers into 16 equal zones (2.5 × 7.5 cm; Fig. 1B). The four zones adjacent to each of the end compartments were designated as the choice areas, while the eight zones in the middle were designated as the neutral area (Fig. 1). Based on the video recordings, we digitized the IDs of zones and the time when focal animals moved into a new zone: we considered a movement a zone border crossing when the tip of the animal’s snout entered a new zone. Subsequently, we calculated the time focal animals spent in the central neutral area and either one of the two choice areas (‘space use by focals’) and counted the total number of zone border crossings by the focal individuals (‘movement activity of focals’). We calculated space use by focals and movement activity of focals both for the first part of trials assessing baseline behaviour and for the second part of trials assessing social and sickness behaviours.

### Statistical analysis

Altogether we performed 55 trials: 28 involving an *Rv*-exposed focal and 27 with an unexposed focal. Due to technical issues, we lost recordings on two trials involving unexposed focals. Furthermore, we excluded 15 trials from the statistical analysis where the infection was not detectable with qPCR in *Rv*-exposed focals and another 13 trials where the *Rv*-exposed stimulus did not appear to be infected (three in trials involving an *Rv*-infected focal and 10 in trials involving a non-infected focal). We also excluded an additional trial where an *Rv*-infected focal individual did not leave the neutral area. This resulted in nine trials where the focal individual was *Rv*-infected and 15 trials where focals were *Rv*-free. The time between *Rv* exposure and the performance of the trials (i.e., 4 *vs.* 8 days) was strongly associated with *Rv* infection intensity (see Results); therefore, we used only the latter as an explanatory variable in our analyses.

Using the data collected in the first part of trials we tested whether focal individuals displayed any preference between the areas when stimulus animals were not yet present. For this, we fitted a Generalized Linear Mixed Model (GLMM) built in the R package *glmmTMB* (Brooks et al. 2017) with negative binomial distribution and logit-link function. In this model ‘space use by focals’ was included as the dependent variable, while ‘*Rv* treatment of focals’ (infected *vs.* unexposed) and ‘type of area’ (choice area next to the infected conspecific *vs*. neutral area *vs*. choice area next to the healthy conspecific) and their two-way interaction were added as fixed factors. As the neutral area was two times larger than the choice areas, we included this information (log-transformed) as an offset into this, and all subsequent models. Trial identity was entered as a random factor, as each focal individual had three data (i.e., the time spent in each of the three areas) in the analysis.

Using the data collected in the second part of the trials, i.e. after the addition of stimulus individuals, we assessed whether focal individuals differentiated between areas within the choice chambers to test for generalized social distancing (Fig. 2A) and avoidance behaviour (Fig. 2B) in healthy juveniles and active self-isolation strategies of infected juveniles (Fig. 2D). For this, we fitted a Linear Mixed Model (LMM) using the *nlme* package (Pinheiro et al. 2023) because negative-binomial models did not yield acceptable model fit (see model assumption testing below). We entered ‘space use by focals’ as the dependent variable and added ‘*Rv* treatment of focals’, ‘type of area’, ‘infection intensity of the stimulus’, and all three- and two-way interactions as fixed factors. As a covariate, we also added ‘space use in the first part of trials’, i.e., the proportion of time a given focal individual spent in the respective area during the first part of the trial, to control for potential side preference. Trial identity was added as a random factor, with three data per focal individual as in the previous model. We square-root transformed ‘space use by focals’ and log-transformed ‘infection intensity of the stimulus’ to enhance the normality of model residuals. To test for generalized social distancing in healthy and self-isolation in infected juveniles we assessed whether focal individuals spent more time in the neutral than the choice areas. To test for avoidance behaviour, we investigated whether focal individuals spent less time in the choice area close to the *Rv*-infected stimulus individuals than close to the non-infected stimulus. Finally, to test if infected focals may have been manipulated by the parasite to stay close to conspecifics in general or to non-infected conspecifics specifically (Fig. 2E), we evaluated whether infected focal individuals spent more time in the choice areas or in the choice areas adjacent to the non-infected stimulus than non-infected focals.

**Figure 2:**
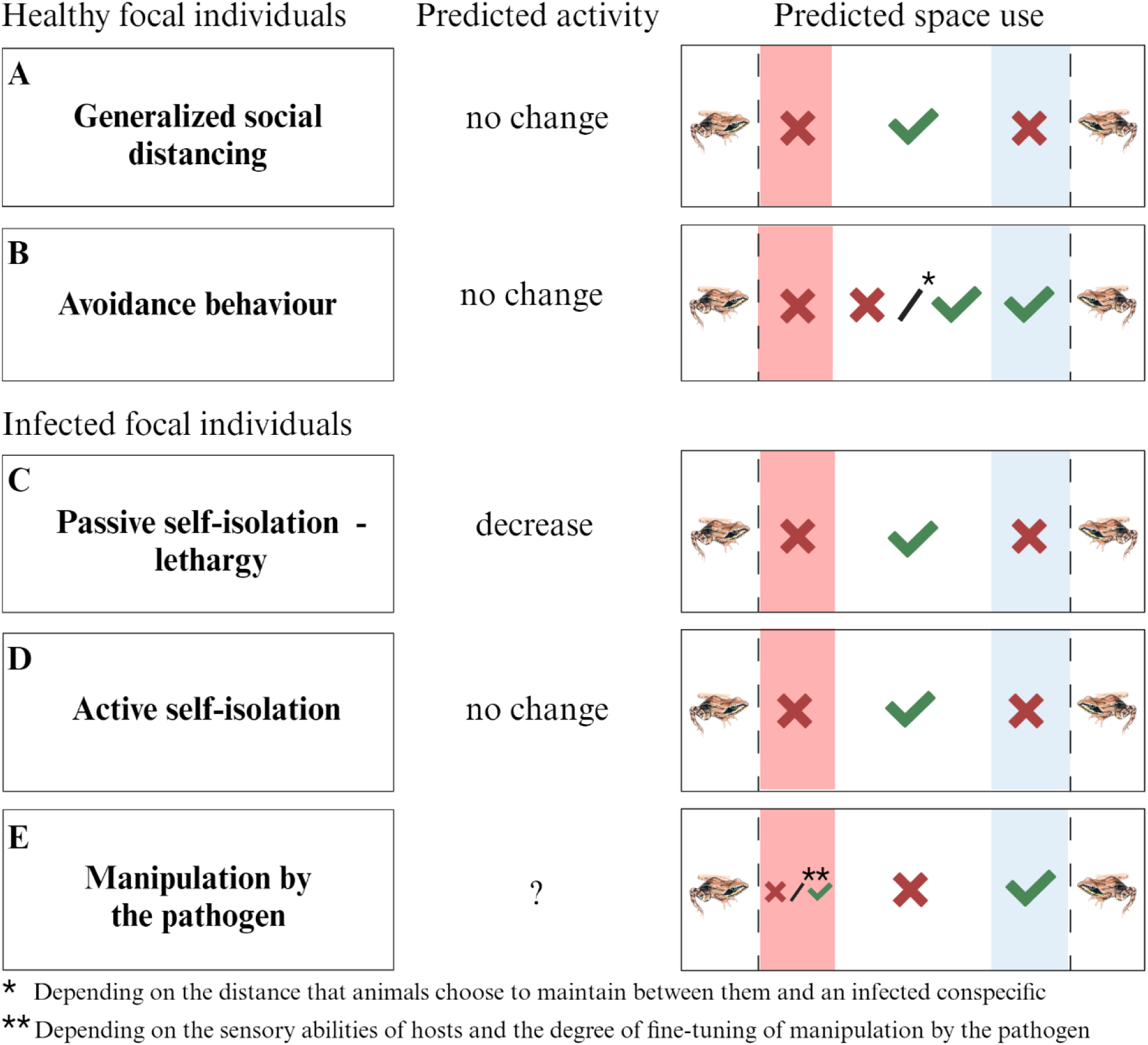
A summary of potential pathogen-induced behaviours of non-infected and infected hosts, along with corresponding predictions that were tested in this study. The red-coloured choice area represents the one adjacent to the infected stimulus while the blue-coloured choice area represents the one adjacent to the non-infected stimulus. Red × symbols denote reduced time relative to the areas marked with green checkmarks.

Furthermore, to test for passive self-isolation *via* lethargy (Fig. 2C), we evaluated whether movement activity differed between infected and healthy focal individuals during the trials. We built a GLMM in *glmmTMB* with negative binomial distribution and logit-link function with ‘movement activity of focals’ as the dependent variable, ‘*Rv* treatment of focals’, ‘type of area’, ‘part of trials’ and all three- and two-way interactions as fixed factors. Trial identity was added as a random factor, as each focal individual had two activity values (one for the first part and one for the second part of the trial).

To test whether the focal individuals’ infection intensity affected their space-use behaviour in the second part of the trials, we used a reduced subset containing the infected focal individuals only. We fitted an LMM with ‘space use by focals’ as the dependent variable, and ‘infection intensity of the focal’, ‘type of area’ and their two-way interaction fitted as fixed effects and ‘space use in the first part of trials’ as a covariate. We added trial identity as a random factor (i.e. three areas per focal individual). Finally, to test for an effect of infection intensity of the focal individuals on their movement activity over the entire trial, we fitted a GLMM with ‘movement activity of the infected focal’ as the dependent variable and ‘type of area’, ‘infection intensity of the focal’, ‘part of trials’ and all three-way and two-way interactions as fixed factors. Trial identity was entered as a random factor, as each focal individual was represented by three data points (i.e., the time spent in each one of the three types of areas) in the analysis.

Error distributions, link functions, and transformations of the depoe applied in the models were chosen after the inspection of residual plots, using the *DHARMa* package for generalized models (Hartig 2020). We applied a backward stepwise model selection procedure to reduce noise in parameter estimates due to the inclusion of non-significant terms (Grafen & Hails 2002; Engqvist 2005). Two variables were never omitted during model reduction regardless of their significance: for space use in the second part of the trials, we always controlled for space use in the first part of the trials; for movement activity, we always retained the treatment effect to which the tested prediction referred. We obtained variance-partitioning statistics for excluded terms by re-entering them to the final model. For these steps, we used type-2 analysis-of-deviance tables (‘Anova’ function of the *car* package). For multiple comparisons derived from the same model to test each prediction, we applied the false discovery rate (FDR) correction method to adjust *P* values (Pike 2011; Lenth et al. 2021). All analyses were performed using R 4.3.2 (R Core Team 2023); our annotated R script is available as supplementary material. Figures 1 and 2 were created with BioRender (BioRender.com).

## RESULTS

We observed no cross-contamination during the experiment: all control juveniles, which had been exposed to the sham treatment, remained *Rv-*free. Amongst the *Rv*-exposed focal animals, infection prevalence was 46.4 % and the median infection intensity in positive individuals was 3.18 × 10^5^ pfu ml^−1^ (min-max: 11 - 6.45 × 10^7^ pfu ml^−1^). In *Rv*-exposed stimulus animals, infection prevalence was 67.9 % with the median infection intensity in positive individuals being 1.04 × 10^5^ pfu ml^−1^ (min-max: 15 - 9.12 × 10^7^ pfu ml^−1^). Ranavirus-positive individuals showed significantly higher viral loads after 8 days post-exposure (median: 2.28 × 10^6^ pfu ml^−1^, min-max 55 - 9.12 × 10^7^ pfu ml^−1^) compared to 4 days post-exposure (median: 5845.5 pfu ml^−1^, min-max 11 - 1.76 × 10^7^ pfu ml^−1^; Welch two sample t-test; t = 3.087, *P* < 0.001). Mortality in froglets not used in the trials was low during the experiment: only 3 out of 115 non-exposed and 12 out of 116 *Rv*-exposed individuals died, with the median time lag between *Rv* exposure and death being 10 days (min-max: 8 - 12 days).

During the first part of trials assessing baseline behaviour, the space use by focal animals deviated significantly from random: juveniles spent more time in the choice area near the end compartment into which we subsequently placed the infected stimulus than both in the neutral area (odds ratio: 2.04 ± 0.594 SE, z = 2.44, *P* = 0.022; Fig. 3A) and in the choice area close to the end compartment into which we later added the non-infected stimulus (odds ratio: 2.46 ± 0.881 SE, z = 2.523, *P* = 0.022; Fig. 3A), while spending similar time in the latter two areas (odds ratio: 1.21 ± 0.43 SE, z = 0.54, *P* = 0.589). Space use by focal animals was not affected by the *Rv* treatment they had been exposed to, or by the two-way interaction between type of area and *Rv* treatment (Table 1; Fig. 3A).

**Figure 3:**
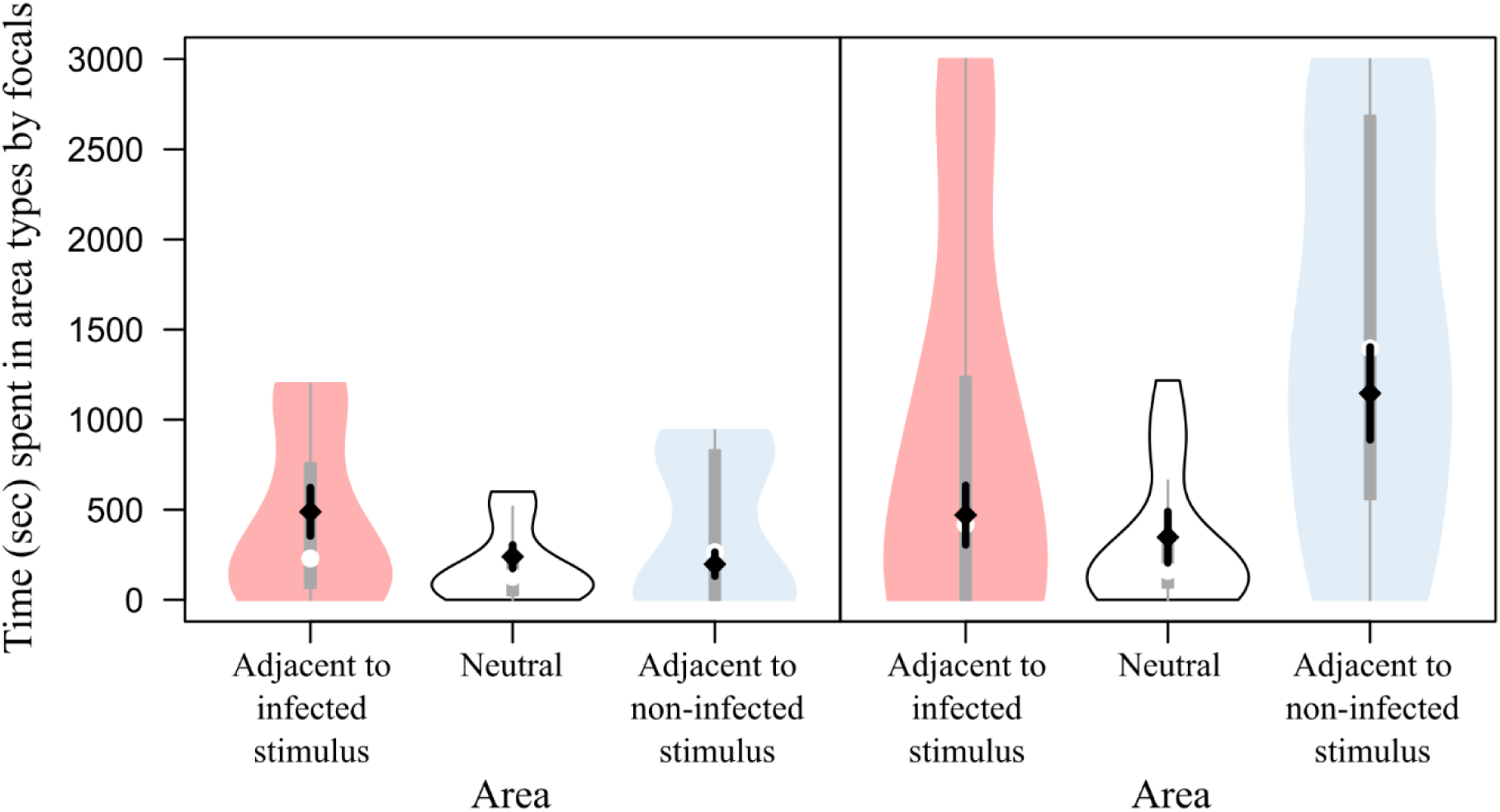
Time spent in the choice areas and the neutral area by focal individuals during the first part of trials assessing baseline behaviour (A) and in the second part of trials assessing social and sickness behaviours (B). In each violin plot, the white dot and the grey box represent the median and the interquartile range, respectively, and the violins are Kernel density plots. The black diamond with whiskers denotes the mean ± standard error estimated from the final statistical models. Note that the second part of trials was two times longer than the first part of trials. Size differences between the choice areas and the neutral area were corrected for unit size.

**Table 1.**
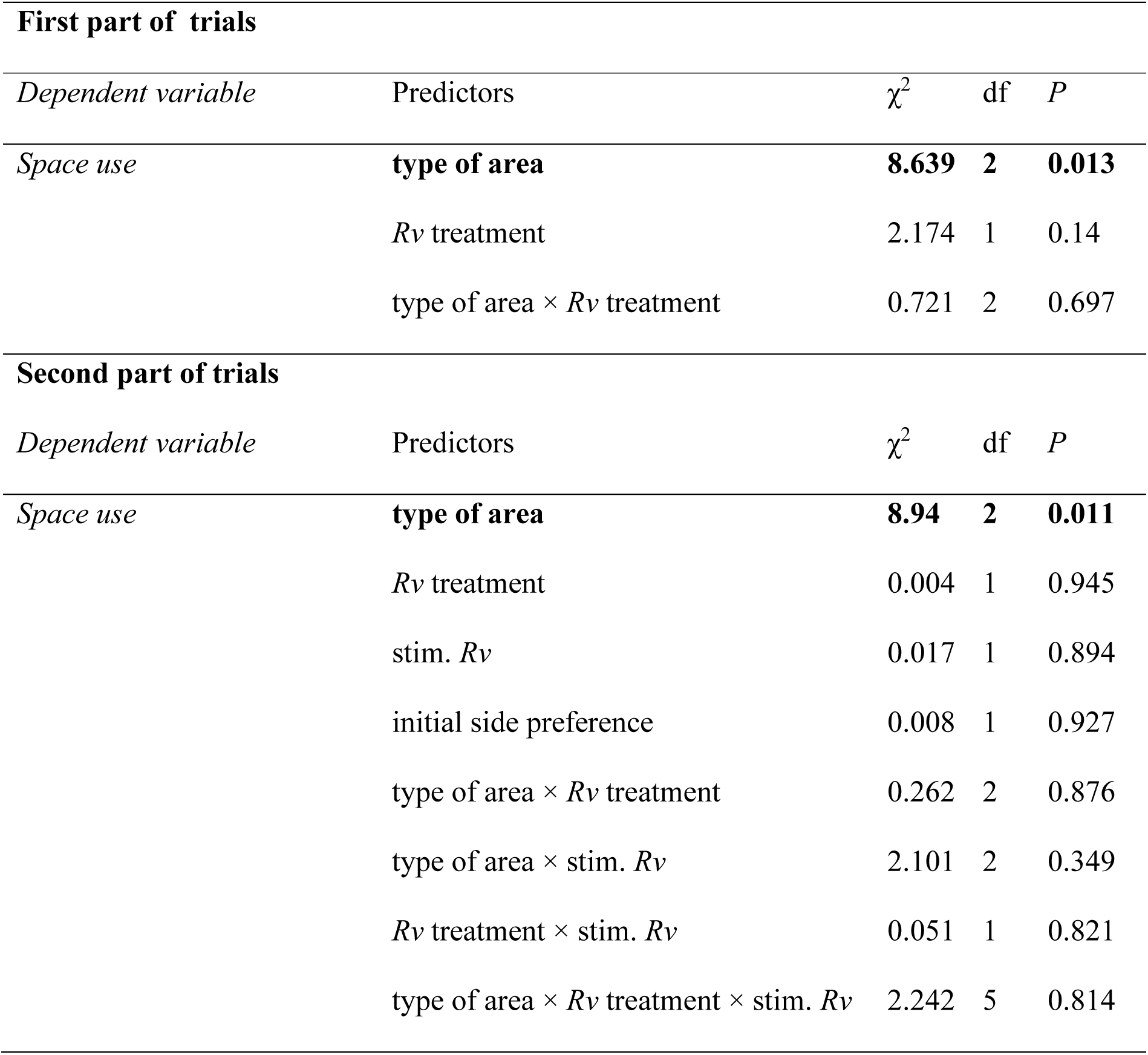
Type-2 analysis-of-deviance tables of the statistical models examining the investigated behavioural components in non-infected and *Ranavirus*-infected focal individuals. Significant effects (*P* < 0.05) are highlighted in bold. Area = two choice areas and the neutral area; *Rv* treatment = infected *vs.* unexposed; stim. *Rv* = log-transformed infection intensity of the stimulus; initial side preference = space use in the first part of trials.

Following the addition of stimulus animals, space use by focal individuals was not affected by their *Rv* treatment, their space use in the first part of trials, or the infection intensity of the stimulus, but was significantly affected by the type of area (Table 1). Specifically, they spent significantly more time in the choice area near the non-infected stimulus animal than near the infected conspecific (t45 = 2.254, *P* = 0.045; Fig. 3B; see avoidance in Fig. 2A). Their time spent in the neutral area was similar to the latter (t45 = 0.55, *P* = 0.582; Fig. 3B) and greater than the former (t45 = 2.80, *P* = 0.022; Fig. 3B). All two-way interactions and the three-way interaction were non-significant (Table 1). The time spent on average in the two choice areas did not differ from the time spent in the neutral area either in non-infected focals (t43 = 1.20, *P* = 0.237; see generalized social distancing in Fig. 2A) or in infected focals (t43 = 1.55, *P* = 0.237; see active self-isolation in Fig. 2D). Compared to the non-infected focal individuals, infected focals did not spend more time near conspecifics (t66 = 0.374, *P* = 0.709) or close to non-infected conspecifics specifically (t66 = 0.301, *P* = 0.764; see manipulation by the pathogen in Fig. 2E). Within infected focal individuals, space use was not influenced by the infection intensity we detected in them after trials, as shown by the non-significant interaction between the type of area and *Rv* load (LMM: χ^2^2 = 4.594, *P* = 0.101).

The movement activity over the entire trial did not differ between infected (3.25 ± 0.77) and non-infected (4.35 ± 0.78) focal individuals (GLMM: χ^2^1 = 1.039, *P* = 0.307; see passive self-isolation in Fig. 2C). Additionally, activity was not affected by the part of the trials (χ^2^1 = 0.128, *P* = 0.72), but the effect of area type was significant (χ^2^2 = 12.781, *P* = 0.002). Specifically, the frequency of zone border crossings was lower in the neutral area (2.49 ± 0.48) than both in the choice area adjacent to the infected stimulus (4.88 ± 0.95; odds ratio: 0.51 ± 0.10 SE, z = 3.31, *P* = 0.003) and in the choice area close to the non-infected stimulus (4.36 ± 0.90; odds ratio: 0.57 ± 0.12 SE, z = 2.69, *P* = 0.011), while it did not differ between the two choice areas (odds ratio: 1.12 ± 0.24 SE, z = 0.53, *P* = 0.598). All two-way interactions and the three-way interaction were non-significant (all *P* > 0.102). Within infected focal individuals, movement activity over the entire trial was not related to their infection intensity (GLMM: χ^2^1 = 2.452, *P* = 0.117).

## DISCUSSION

The most salient finding of this study is the avoidance behaviour displayed by juvenile agile frogs. Focal individuals avoided the area close to infected conspecifics. Contrary to our expectations, this avoidance behaviour was not affected by the infection status of the focal individual: non-infected and infected juveniles both decreased spatial proximity to infected stimuli to a similar extent. Finally, we did not find indications of any other behavioural change that could be attributed to the presence of *Rv*: non-infected juveniles did not display generalized social distancing and infected focal individuals did not show signs of passive or active self-isolation.

Our results demonstrate that susceptible juvenile amphibians can display avoidance behaviour when encountering *Rv*-infected conspecifics. This implies that already as recently metamorphosed juveniles, completely naïve to ranavirosis, they can recognize if a conspecific is infected with *Rv*, upon which they discriminate against them by avoiding close spatial association. In the first part of trials, when only focal individuals, but no stimulus individuals were present in choice chambers, focals avoided the neutral area and the choice area close to the non-infected stimulus and showed a preference for the end compartment into which we subsequently placed the infected stimulus. Because end compartments were assigned to treatments randomly we consider this pattern to have resulted from random noise. In contrast, following the appearance of the stimuli, the previous spatial pattern was reversed, which further underlines the presence of strong avoidance behaviour in focal individuals. A recent study concerned with the *R. sylvatica* – *Rv* host-pathogen system came to similar conclusions in the sense that healthy juveniles kept greater distances from infected conspecifics in paired trials performed in an open arena (Le Sage et al. 2022). However, their experimental design, i.e. observation dyads of one infected and one healthy frog moving around freely in the same space, did not allow for determining to what extent the distance between individuals accorded to the preference of the non-infected frog and resulted from its consequent behaviour. By studying the *R. dalmatina* – *Rv* host-pathogen system in choice chambers, we could confirm the presence of avoidance behaviour in healthy juveniles: the increased distance between the infected stimulus and the non-infected focal individuals was not a by-product of generalized social distancing by focals, nor did it result from social distancing by infected stimulus individuals. Avoidance behaviour has also been demonstrated in larvae of the bullfrog *Lithobates catesbeianus*: Kiesecker and colleagues (1999) showed that in choice chambers larvae spatially avoid conspecifics infected with the pathogenic fungus *Candida humicola*. Based on our results and those of the above two studies on amphibians (Kiesecker et al. 1999; Le Sage et al. 2022), we suggest that avoidance behaviour may be a widespread strategy of disease prevention in amphibians, or at least in anurans. This possibility is strengthened by a large number of demonstrations of avoidance behaviour in several other taxa, including fishes (Mikheev et al. 2013; Stephenson et al. 2018; Stephenson 2019), bats (Langwig et al. 2012; Wilcox et al. 2014), primates (Poirotte et al. 2017; Sarabian et al. 2017) and humans (Curtis 2014) and by some more sporadic observations in others, such as in insects (Heinze & Walter 2010), crustaceans (Behringer et al. 2006; Childress et al. 2015) and birds (Zylberberg et al. 2013). Whether the occurrence of avoidance behaviour is even more universal in the animal kingdom than we currently think, or what extrinsic or intrinsic ecological factors limit its evolution remains to be uncovered.

Contrary to our expectations and conclusions of a similar study (Kiesecker et al. 1999), we observed that *Rv*-infected agile frog juveniles spent less time near infected conspecifics than near non-infected conspecifics. The proximate or ultimate mechanisms behind this behaviour are far from self-explanatory. As a first line of defence, avoidance behaviour is generally thought to evolve to lower the threat of becoming infected (Romano et al. 2020). Accordingly, non-infected individuals have been documented to show avoidance behaviours in a diverse set of taxa (Townsend et al. 2020; Stockmaier et al. 2021), but reports on similar behaviours in already infected individuals are, to our knowledge, non-existent. The question arises as to why infected individuals may show avoidance behaviour, even though they have already contracted the disease. There are at least three plausible explanations. First, the drive underlying avoidance behaviour may be hard-wired so that becoming infected does not significantly weaken it. Second, infected individuals may have been in a too early stage of their infection for ranavirosis to be able to induce detectable changes in their behaviour (Jancovich et al. 1997). This hypothesis is supported by the fact that severe ranavirosis is frequently accompanied by clinical symptoms such as systemic haemorrhages, ulcerations of the skin, and lethargy (Gray et al. 2009), none of which we observed in our study animals during the assays. Third, avoidance behaviour may be adaptive not only for non-infected individuals but for already infected ones as well: by avoiding infected conspecifics, they may lower the chance of re-infection with the same pathogen or with new ones. Higher doses of infection come with increased virulence and decreased average survival time in nature (Anderson & May 1978) and this is also true for larval agile frogs (Herczeg et al. 2023). Further, by staying away from diseased conspecifics, infected individuals may avoid infection with additional new pathogens which may be more likely carried by individuals (co-)infected with *Rv*, and thereby evade potentially disastrous outcomes of co-infection (Herczeg et al. 2021).

In our study system, in the case of infected focal individuals, the observed increase in spatial proximity to the healthy conspecific might not necessarily result from avoidance of the infected stimulus but may be the manifestation of the pathogen manipulating the hosts’ behaviour: infection with *Rv* may have driven froglets to stay close to non-infected conspecifics. Such manipulation by the pathogen to facilitate its transmission is well-known from many host-pathogen systems (Klein 2003; Hernandez-Caballero et al. 2022). Besides other pathogens, viruses are also known to be able to modify the behaviour of their hosts to their advantage (Goulson 1997; Hanlon 2013). However, the pervasive presence of manipulation by the pathogen in our study system is rendered unlikely by the observation that infected focals did not spend more time near conspecifics than in the neutral area, nor did they spend more time near the non-infected conspecifics than did non-infected focals.

We found no sign of the presence of either passive or active self-isolation in *Rv*-infected froglets. However, passive and active self-isolation is often difficult to discern given that side effects of infection and active, adaptive host responses are intertwined (Poulin 1995; Stockmaier et al. 2021). Self-isolation behaviour of infected hosts (Shattuck & Muehlenbein 2015) can occur passively *via* disease-caused lethargy (Van Kerckhove et al. 2013; Hammerstein & Noë 2016; Lopes et al. 2016). Here, we found no evidence of lethargy because movement activity did not differ between *Rv*-infected and non-infected focals. This aligns with the results of previous studies on pathogen avoidance behaviour in amphibians, which also did not document lethargy in experimental animals (Kiesecker et al. 1999; Le Sage et al. 2022). The discrepancy between pathological descriptions and behavioural experiments may be because apathy does appear in animals suffering from ranavirosis but only in the terminal stage of disease progression (Jancovich et al. 1997), while experimenters aim to use animals that still show ‘normal’ behaviours. However, even if passive self-isolation is lacking, infected individuals may still express active self-isolation, but this behavioural strategy is thought to evolve only in social animals *via* kin selection and has been documented solely in eusocial insects and humans (Scott & Duncan 2001; Heinze & Walter 2010; Bos et al. 2012; Stroeymeyt et al. 2018). The absence of active self-isolation in the non-social juvenile agile frogs accords with this theory.

Surprisingly, the behaviour of focal animals did not seem to be influenced by *Rv* infection intensity measured in stimulus or in focal individuals. Le Sage and colleagues (2022) found that the distance between *Rv*-infected and non-infected frogs increased with higher *Rv* infection intensity, while Kiesecker et al. (1999) did not report on pathogen loads. The lack of an effect of infection intensity in our study is, however, in congruence with the observation that infected and non-infected animals did not differ in any of the quantified behaviours. It is also worth noting that only nine individuals were *Rv* positive and the effect of infection intensity may have surfaced at larger sample sizes.

In conclusion, our results provide evidence for the presence of avoidance behaviour in juvenile agile frogs, and this behaviour was not only expressed by non-infected but also by *Rv*-infected froglets. At the same time, behavioural patterns hinting towards the existence of generalized social distancing in non-infected individuals or self-isolation in infected juveniles were absent. A better understanding of the behaviour of hosts in the presence of pathogens as well as of the cues they use to recognize contagious individuals will be of vital importance for devising effective mitigation methods, while such knowledge can also help parameterise epidemiological models and thereby enhance the precision of theoretical predictions, ultimately resulting in more effective mitigation.

## ACKNOWLEDGEMENT

We thank M. Szederkényi, and A. Kraxner for field and technical assistance during the experiment. The Frog Virus 3 isolate was obtained from R.E. Marschang (Laboklin GmbH). We thank A. Doszpoly (HUN-REN ÁOTKI) for help with Ranavirus maintenance. Also, we thank the members of the Department of Plant Pathology (HUN-REN ATK) for allowing us to use their facility during laboratory work. We thank B. Bombay for the agile frog painting. The study was funded by the National Research, Development and Innovation Office of Hungary (NKFIH, grants K-124375 and K-147500 for A. Hettyey). G. Horváth gained support from the postdoctoral research grant of the National Research, Development and Innovation Fund (NKFIH, PD 132041) and A. Hettyey from the János Bolyai Research Scholarship of the Hungarian Academy of Sciences (MTA, BO/00667/21/8). The authors were supported by the New National Excellence Program of the Ministry for Innovation and Technology of the National Research, Development and Innovation Fund (ÚNKP-21-3 to A. Kásler, ÚNKP-19-4 and ÚNKP-22-5 to A. Hettyey, ÚNKP-23-4 to J. Ujszegi). This project received funding from the HUN-REN Hungarian Research Network. The first author dedicated this work to the loving memory of László Hajcsák.

## DATA ACCESSIBILITY

Data is available from Figshare Digital Repository doi: 10.6084/m9.figshare.25568805. The DOI becomes active upon acceptance of the manuscript.

## ETHICS STATEMENT

All experimental procedures were approved by the Ethical Commission of the HUN-REN ATK NÖVI under Good Scientific Practice guidelines and national legislation and we carried out experiments according to permits issued by the Government Agency of Pest County (Department of Environmental Protection and Nature Conservation, PE/EA/295-7/2018 and PE/EA/58-4/2019).

## AUTHORS CONTRIBUTIONS

Dávid Herczeg and Attila Hettyey conceived the study and designed the methodology; Dávid Herczeg, Andrea Kásler, Dóra Holly, Veronika Bókony, Nikolett Ujhegyi, Tibor Papp, János Ujszegi and Attila Hettyey collected the data; Gergely Horváth, Veronika Bókony and Dávid Herczeg analysed the data; Dávid Herczeg, Gábor Herczeg and Attila Hettyey led the writing of the manuscript. All authors contributed critically to the drafts and gave final approval for publication.

## REFERENCES

Amoroso, C.R., Kappeler, P.M., Fichtel, C. & Nunn, C.L. (2019) Fecal contamination, parasite risk, and waterhole use by wild animals in a dry deciduous forest. Behavioral Ecology and Sociobiology, 73, 153.10.1007/s00265-019-2769-6

Anderson, R.M. & May, R.M. (1978) Regulation and Stability of Host-Parasite Population Interactions: I. Regulatory Processes. Journal of Animal Ecology, 47, 219–247.10.2307/3933

Behringer, D.C., Butler, M.J. & Shields, J.D. (2006) Avoidance of disease by social lobsters. Nature, 441, 421–421.10.1038/441421a

Behringer, D.C., Karvonen, A. & Bojko, J. (2018) Parasite avoidance behaviours in aquatic environments. Philosophical Transactions of the Royal Society B: Biological Sciences, 373, 20170202.doi:10.1098/rstb.2017.0202

Bos, N., Lefévre, T., Jensen, A.B. & D’Ettorre, P. (2012) Sick ants become unsociable. Journal of Evolutionary Biology, 25, 342–351.10.1111/j.1420-9101.2011.02425.x

Brooks, M.E., Kristensen, K., van Benthem, K.J., Magnusson, A., Berg, C.W., Nielsen, A., Skaug, H.J., Mächler, M. & Bolker, B.M. (2017) glmmTMB balances speed and flexibility among packages for zero-inflated generalized linear mixed modeling. The R Journal, 9, 378–400

Brunner, J.L., Storfer, A., Gray, M.J. & Hoverman, J.T. (2015) Ranavirus Ecology and Evolution: From Epidemiology to Extinction. Ranaviruses: Lethal Pathogens of Ectothermic Vertebrates (eds M.J. Gray & V.G. Chinchar), pp. 71-104. Springer International Publishing, Cham.10.1007/978-3-319-13755-1_4

Childress, M.J., Heldt, K.A. & Miller, S.D. (2015) Are juvenile Caribbean spiny lobsters (Panulirus argus) becoming less social? ICES Journal of Marine Science, 72, i170–i176.10.1093/icesjms/fsv045

Collins, J.P. (2010) Amphibian decline and extinction: what we know and what we need to learn. Diseases of aquatic organisms, 92, 93–99.10.3354/dao02307

Curtis, V.A. (2014) Infection-avoidance behaviour in humans and other animals. Trends in Immunology, 35, 457–464.10.1016/j.it.2014.08.006

Duffus, A.L.J., Waltzek, T.B., Stöhr, A.C., Allender, M.C., Gotesman, M., Whittington, R.J., Hick, P., Hines, M.K. & Marschang, R.E. (2015) Distribution and Host Range of Ranaviruses. Ranaviruses: Lethal Pathogens of Ectothermic Vertebrates (eds M.J. Gray & V.G. Chinchar), pp. 9-57. Springer International Publishing, Cham.10.1007/978-3-319-13755-1_2

Earl, J.E., Chaney, J.C., Sutton, W.B., Lillard, C.E., Kouba, A.J., Langhorne, C., Krebs, J., Wilkes, R.P., Hill, R.D., . . . Gray, M.J. (2016) Ranavirus could facilitate local extinction of rare amphibian species. Oecologia, 182, 611–623.10.1007/s00442-016-3682-6

Engqvist, L. (2005) The mistreatment of covariate interaction terms in linear model analyses of behavioural and evolutionary ecology studies. Animal Behaviour, 70, 967–971.10.1016/j.anbehav.2005.01.016

Fisher, M.C. & Garner, T.W.J. (2020) Chytrid fungi and global amphibian declines. Nature Reviews Microbiology, 18, 332–343.10.1038/s41579-020-0335-x

Friard, O. & Gamba, M. (2016) BORIS: a free, versatile open-source event-logging software for video/audio coding and live observations. Methods in Ecology and Evolution, 7, 1325–1330.10.1111/2041-210X.12584

Geffre, A.C., Gernat, T., Harwood, G.P., Jones, B.M., Morselli Gysi, D., Hamilton, A.R., Bonning, B.C., Toth, A.L., Robinson, G.E., . . . Dolezal, A.G. (2020) Honey bee virus causes context-dependent changes in host social behavior. Proceedings of the National Academy of Sciences, 117, 10406–10413.10.1073/pnas.2002268117

Gosner, K.L. (1960) A simplified table for staging anuran embryos and larvae with notes on identification. Herpetologica, 16, 183–190

Goulson, D. (1997) Wipfelkrankheit: modification of host behaviour during baculoviral infection. Oecologia, 109, 219–228.10.1007/s004420050076

Grafen, A. & Hails, R. (2002) Modern statistics for the life sciences. Oxford University Press, Oxford.

Gray, M.J., Miller, D.L. & Hoverman, J.T. (2009) Ecology and pathology of amphibian ranaviruses. Diseases of aquatic organisms, 87, 243–266243.10.3354/dao02138

Hafer, N. (2016) Conflicts over host manipulation between different parasites and pathogens: Investigating the ecological and medical consequences. BioEssays, 38, 1027–1037.10.1002/bies.201600060

Hammerstein, P. & Noë, R. (2016) Biological trade and markets. Philosophical Transactions of the Royal Society B: Biological Sciences, 371, 20150101.10.1098/rstb.2015.0101

Hanlon, C.A. (2013) Chapter 5 - Rabies in Terrestrial Animals. Rabies (Third Edition) (ed. A.C. Jackson), pp. 179–213. Academic Press, Boston.10.1016/B978-0-12-396547-9.00005-5

Hartig, F. (2020) DHARMa: residual diagnostics for hierachical (multi-level/mixed) regression models. R package version 0.3.3.0.

Heinze, J. & Walter, B. (2010) Moribund Ants Leave Their Nests to Die in Social Isolation. Current Biology, 20, 249–252.10.1016/j.cub.2009.12.031

Herczeg, D., Holly, D., Kásler, A., Bókony, V., Papp, T., Takács-Vágó, H., Ujszegi, J. & Hettyey, A. (2023) Amphibian larvae benefit from a warm environment under simultaneous threat from chytridiomycosis and ranavirosis. Oikos, 2023, e09953.10.1111/oik.09953

Herczeg, D., Ujszegi, J., Kásler, A., Holly, D. & Hettyey, A. (2021) Host–multiparasite interactions in amphibians: a review. Parasites & Vectors, 14, 296. 10.1186/s13071-021-04796-1

Hernandez-Caballero, I., Garcia-Longoria, L., Gomez-Mestre, I. & Marzal, A. (2022) The Adaptive Host Manipulation Hypothesis: Parasites Modify the Behaviour, Morphology, and Physiology of Amphibians. Diversity.

Jancovich, J.K., Davidson, E.W., Morado, J.F., Jacobs, B.L. & Collins, J.P. (1997) Isolation of a lethal virus from the endangered tiger salamander *Ambystoma tigrinum stebbinsi*. Diseases of aquatic organisms, 31, 161–167

Kazlauskas, N., Klappenbach, M., Depino, A.M. & Locatelli, F.F. (2016) Sickness Behavior in Honey Bees. Frontiers in Physiology, 7.10.3389/fphys.2016.00261

Kiesecker, J.M. & Skelly, D.K. (2000) Choice of Oviposition Site by Gray Treefrogs: The Role of Potential Parasitic Infection. Ecology, 81, 2939–2943.10.2307/177354

Kiesecker, J.M., Skelly, D.K., Beard, K.H. & Preisser, E. (1999) Behavioral reduction of infection risk. Proceedings of the National Academy of Sciences, 96, 9165–9168.10.1073/pnas.96.16.9165

Kik, M., Martel, A., Sluijs, A.S.-v.d., Pasmans, F., Wohlsein, P., Gröne, A. & Rijks, J.M. (2011) Ranavirus-associated mass mortality in wild amphibians, the Netherlands, 2010: a first report. Veterinary journal, 190, 284–286. 10.1016/j.tvjl.2011.08.031

Klein, S.L. (2003) Parasite manipulation of the proximate mechanisms that mediate social behavior in vertebrates. Physiology & Behavior, 79, 441–449.10.1016/S0031-9384(03)00163-X

Landsberg, J.H., Kiryu, Y., Tabuchi, M., Waltzek, T.B., Enge, K.M., Reintjes-Tolen, S., Preston, A. & Pessier, A.P. (2013) Co-infection by alveolate parasites and frog virus 3- like ranavirus during an amphibian larval mortality event in Florida, USA. Diseases of aquatic organisms, 105 2, 89–99.10.3354/dao02625

Langwig, K.E., Frick, W.F., Bried, J.T., Hicks, A.C., Kunz, T.H. & Marm Kilpatrick, A. (2012) Sociality, density-dependence and microclimates determine the persistence of populations suffering from a novel fungal disease, white-nose syndrome. Ecology Letters, 15, 1050–1057.10.1111/j.1461-0248.2012.01829.x

Le Sage, E.H., Diamond, M. & Crespi, E.J. (2022) Ranavirus infection-induced avoidance behaviour in wood frog juveniles: do amphibians socially distance? Biology Letters, 18, 20220359.10.1098/rsbl.2022.0359

Lenth, R.V., Buerkner, P., Herve, M., Love, J., Riebl, H. & Singmann, H. (2021) emmeans: Estimated marginal means, aka least-squares means. CRAN.

Lopes, P.C., Block, P. & König, B. (2016) Infection-induced behavioural changes reduce connectivity and the potential for disease spread in wild mice contact networks. Scientific Reports, 6, 31790.10.1038/srep31790

Luedtke, J.A., Chanson, J., Neam, K., Hobin, L., Maciel, A.O., Catenazzi, A., Borzée, A., Hamidy, A., Aowphol, A., . . . Stuart, S.N. (2023) Ongoing declines for the world’s amphibians in the face of emerging threats. Nature, 622, 308–314.10.1038/s41586-023-06578-4

McMahon, T.A., Hill, M.N., Lentz, G.C., Scott, E.F., Tenouri, N.F. & Rohr, J.R. (2021) Amphibian species vary in their learned avoidance response to the deadly fungal pathogen *Batrachochytrium dendrobatidis*. Journal of Applied Ecology, 58, 1613–1620.10.1111/1365-2664.13932

McMahon, T.A., Sears, B.F., Venesky, M.D., Bessler, S.M., Brown, J.M., Deutsch, K., Halstead, N.T., Lentz, G., Tenouri, N., . . . Rohr, J.R. (2014) Amphibians acquire resistance to live and dead fungus overcoming fungal immunosuppression. Nature, 511, 224–227.10.1038/nature13491

Mikheev, V.N., Pasternak, A.F., Taskinen, J. & Valtonen, T.E. (2013) Grouping facilitates avoidance of parasites by fish. Parasites & Vectors, 6, 301.10.1186/1756-3305-6-301

Moleón, M., Martínez-Carrasco, C., Muellerklein, O.C., Getz, W.M., Muñoz-Lozano, C. & Sánchez-Zapata, J.A. (2017) Carnivore carcasses are avoided by carnivores. Journal of Animal Ecology, 86, 1179–1191.10.1111/1365-2656.12714

Paciência, F.M.D., Rushmore, J., Chuma, I.S., Lipende, I.F., Caillaud, D., Knauf, S. & Zinner, D. (2019) Mating avoidance in female olive baboons (*Papio anubis*) infected by Treponema pallidum. Science Advances, 5, eaaw9724.10.1126/sciadv.aaw9724

Pike, N. (2011) Using false discovery rates for multiple comparisons in ecology and evolution. Methods in Ecology and Evolution, 2, 278–282.10.1111/j.2041-210X.2010.00061.x

Pinheiro, J., Bates, D. & Team, R.C. (2023) nlme: Linear and Nonlinear Mixed Effects Models. R package version 3.1–163. CRAN.

Poirotte, C., Massol, F., Herbert, A., Willaume, E., Bomo, P.M., Kappeler, P.M. & Charpentier, M.J.E. (2017) Mandrills use olfaction to socially avoid parasitized conspecifics. Science Advances, 3, e1601721.10.1126/sciadv.1601721

Poulin, R. (1995) “Adaptive” changes in the behaviour of parasitized animals: A critical review. International Journal for Parasitology, 25, 1371–1383.10.1016/0020-7519(95)00100-X

Price, Stephen J., Garner, Trenton W.J., Nichols, Richard A., Balloux, F., Ayres, C., Mora-Cabello de Alba, A. & Bosch, J. (2014) Collapse of amphibian communities due to an introduced Ranavirus. Current Biology, 24, 2586–2591.10.1016/j.cub.2014.09.028

R Core Team (2023) R: A language and environment for statistical computing. R Foundation for Statistical Computing, Vienna, Austria.

Romano, V., MacIntosh, A.J.J. & Sueur, C. (2020) Stemming the Flow: Information, Infection, and Social Evolution. Trends in Ecology & Evolution, 35, 849–853.10.1016/j.tree.2020.07.004

Sarabian, C., Curtis, V. & McMullan, R. (2018) Evolution of pathogen and parasite avoidance behaviours. Philos Trans R Soc Lond B Biol Sci, 373.10.1098/rstb.2017.0256

Sarabian, C., Ngoubangoye, B. & MacIntosh, A.J.J. (2017) Avoidance of biological contaminants through sight, smell and touch in chimpanzees. Royal Society Open Science, 4, 170968.10.1098/rsos.170968

Scheele, B.C., Pasmans, F., Skerratt Lee, F., Berger, L., Martel, A., Beukema, W., Acevedo Aldemar, A., Burrowes Patricia, A., Carvalho, T., Canessa, S. (2019) Amphibian fungal panzootic causes catastrophic and ongoing loss of biodiversity. Science, 363, 1459–1463.10.1126/science.aav0379

Schock, D.M., Bollinger, T.K., Gregory Chinchar, V., Jancovich, J.K. & Collins, J.P. (2008) Experimental evidence that amphibian ranaviruses are multi-host pathogens. Copeia, 2008, 133–143.10.1643/cp-06-134

Scott, S. & Duncan, C.J. (2001) Biology of Plagues: Evidence from Historical Populations. Cambridge University Press, Cambridge. DOI: 10.1017/CBO9780511542527

Shattuck, E.C. & Muehlenbein, M.P. (2015) Human sickness behavior: Ultimate and proximate explanations. American Journal of Physical Anthropology, 157, 1–18.10.1002/ajpa.22698

Shine, R., Amiel, J., Munn, A.J., Stewart, M., Vyssotski, A.L. & Lesku, J.A. (2015) Is “cooling then freezing” a humane way to kill amphibians and reptiles? Biology Open, 4, 760–763.10.1242/bio.012179

Stephenson, J.F. (2019) Parasite-induced plasticity in host social behaviour depends on sex and susceptibility. Biology Letters, 15, 20190557.10.1098/rsbl.2019.0557

Stephenson, J.F., Perkins, S.E. & Cable, J. (2018) Transmission risk predicts avoidance of infected conspecifics in Trinidadian guppies. Journal of Animal Ecology, 87, 1525–1533.10.1111/1365-2656.12885

Stilwell, N.K., Whittington, R.J., Hick, P.M., Becker, J.A., Ariel, E., van Beurden, S., Vendramin, N., Olesen, N.J. & Waltzek, T.B. (2018) Partial validation of a TaqMan real-time quantitative PCR for the detection of ranaviruses. Diseases of aquatic organisms, 128, 105–116.10.3354/dao03214

Stockmaier, S., Bolnick, D.I., Page, R.A. & Carter, G.G. (2020) Sickness effects on social interactions depend on the type of behaviour and relationship. Journal of Animal Ecology, 89, 1387–1394.10.1111/1365-2656.13193

Stockmaier, S., Stroeymeyt, N., Shattuck, E.C., Hawley, D.M., Meyers, L.A. & Bolnick, D.I. (2021) Infectious diseases and social distancing in nature. Science, 371, eabc8881.10.1126/science.abc8881

Stroeymeyt, N., Casillas-Pérez, B. & Cremer, S. (2014) Organisational immunity in social insects. Current Opinion in Insect Science, 5, 1–15.10.1016/j.cois.2014.09.001

Stroeymeyt, N., Grasse, A.V., Crespi, A., Mersch, D.P., Cremer, S. & Keller, L. (2018) Social network plasticity decreases disease transmission in a eusocial insect. Science, 362, 941–945.10.1126/science.aat4793

Townsend, A.K., Hawley, D.M., Stephenson, J.F. & Williams, K.E.G. (2020) Emerging infectious disease and the challenges of social distancing in human and non-human animals. Proceedings of the Royal Society B: Biological Sciences, 287, 20201039.10.1098/rspb.2020.1039

USEPA (2002) Methods for measuring the acute toxicity of effluents and receiving waters to freshwater and marine organisms [internet]. United States Environmental Protection Agency Office of Water

Van Kerckhove, K., Hens, N., Edmunds, W.J. & Eames, K.T.D. (2013) The Impact of Illness on Social Networks: Implications for Transmission and Control of Influenza. American Journal of Epidemiology, 178, 1655–1662.10.1093/aje/kwt196

Wake, D.B. & Vredenburg, V.T. (2008) Are we in the midst of the sixth mass extinction? A view from the world of amphibians. Proceedings of the National Academy of Sciences, 105, 11466–11473.10.1073/pnas.0801921105

Weinstein, S.B., Buck, J.C. & Young, H.S. (2018) A landscape of disgust. Science, 359, 1213–1214.doi:10.1126/science.aas8694

Wilcox, A., Warnecke, L., Turner, J.M., McGuire, L.P., Jameson, J.W., Misra, V., Bollinger, T.C. & Willis, C.K.R. (2014) Behaviour of hibernating little brown bats experimentally inoculated with the pathogen that causes white-nose syndrome. Animal Behaviour, 88, 157–164.10.1016/j.anbehav.2013.11.026

Zylberberg, M., Klasing, K.C. & Hahn, T.P. (2013) House finches (*Carpodacus mexicanus*) balance investment in behavioural and immunological defences against pathogens. Biology Letters, 9, 20120856.10.1098/rsbl.2012.0856

